# Development and Characterization of a New Endoscopic Drug Eluting Platform with Proven Efficacy in Acute and Chronic Experimental Colitis

**DOI:** 10.1101/712224

**Authors:** I Bon, M Cano-Sarabia, N de la Ossa, R Bartolí, V Lorenzo-Zúñiga

## Abstract

**Background&Aims:** Mucosal lesions refractory to biological treatments represent unmet needs in patients with inflammatory bowel disease (IBD) that require new treatment modalities. We developed and characterized a new endoscopic drug-eluting hydrogel (CoverGel) with proven efficacy in acute and chronic experimental colitis (EC) in rats

**Methods:** CoverGel was developed based on appropriate rheological, drug release, gelation, structural and degradation properties capacities to allow endoscopic application. Experimental colitis (EC) was induced by TNBS application in rats. In acute EC 40 rats were randomized in 5 groups (8 each): sham, control, CoverGel, CoverGel+Infliximab (IFX) and CoverGel+Vedolizumab (VDZ). In chronic EC 12 rats were randomized in 2 groups (6 each): IFX s.c and CoverGel+IFX. Endoscopic, histological and blood test were performed during follow-up to evaluate clinical success. Antibodies to IFX (ATIs) were evaluated in chronic EC animal study

**Results:** CoverGel is a biocompatible and bioadhesive reverse thermo sensitive gelation hydrogel with macroporous structure and drug release capacity. In acute EC animals treated with CoverGel+IFX or CoverGel+VDZ showed significantly clinical success (weight recovery, mucosal restoration and bacterial translocation) as compare with controls and animals without bioactive drug. In chronic EC animal study, clinical efficacy was comparable in both groups. Levels of ATIs were significantly lower in animals treated with CoverGel+IFX vs. IFX s.c (0.90 ± 0.06 μg/mL-c vs. 1.97 ± 0.66 μg/mL-c, p=0.0025).

**Conclusions:** CoverGel is an endoscopic vehicle to locally deliver biological drugs with proven efficacy in acute and chronic EC in rats and inducing less immunogenicity reaction.

## INTRODUCTION

Interventional inflammatory bowel disease (IBD) endoscopy has an expanding role in the era of biologic therapy. A significant number of patients have mucosal lesions refractory to biological treatments or have disease-associated adverse events that require surgical intervention such as strictures or neoplasia, despite the availability and growing use of anti-tumor necrosis factor (TNF), anti-integrin, anti-interleukin and new small-molecule^1,2^. These drugs have changed the way of treating IBD patients, but those situations represent unmet needs that require new treatment modalities to prolong the intervention-free period. Endoscopic shielding technique with hydrogels has been proposed by our group as a less invasive approach to reduce or postpone the need for surgical resection^3,4^.

Hydrogels or hydrophilic gels are cross-linked polymeric three-dimensional networks which present soft tissue-like elastic, non-toxic and biodegradable properties while being responsive to stimulus^5–8^. Nowadays we do not have any valid endoscopic treatment to IBD mucosal lesions in patients with refractory or loss of response to biological therapy. The purpose of the present study was to develop and characterize a new endoscopic drug-eluting platform and test its efficacy in acute and chronic experimental colitis in rats.

## METHODS

### Characterization of the drug-eluting platform

We have developed a new hydrogel (CoverGel) comprising specific amounts of hyaluronic acid (HA), methylcellulose (MC) and Poloxamer Pluronic-F127 (F127). This platform has been patented (PCT/EP2017/068876). The composition of CoverGel is based on the combination of different substances to create this solution, which can be prepared before the procedure.

Our goal was to improve the beneficial effects of HA onto the colonic mucosa by adding different substances to increase biocompatibility, bioadhesibility and bioactive properties, and make it suitable to be administered through the endoscope’s working channel. For this, we evaluated different aspects: rheological properties (viscosity, adherence and microstructure), gelation, degradation, drug absorbance, drug release and biocompatibility. Four different original combinations (Colon dressing, CD) prototypes of materials (CD1, CD2, CD3, CD4) were tested for viscosity, adhesion force and microstructure. The best composition for the drug-eluting platform was different percentages of HA, MC and PF127. All compounds of the product are well known and have proved safety. The composition of the invention is biodegradable, is easy to administrate, remains adhered to the mucosal lesion and is a drug-release composition.

#### Rheological test

(i) Viscosity was evaluated in a rheometer Haake RheoStress (Thermo Fisher Scientific, Waltham, MA USA) with a C60/1°Ti probe and a gap set of 0.053 mm. Rotation ramp went from 0–300 s–1 in 30 seconds. For each study, Viscosity (η) was measured as a function of shear rate (γ) at 22°C (room temperature) and 37°C (body temperature), (ii) Adhesion was measured with a Texture Analyzer TA.XT Plus (Stable Micro Systems, Surrey, UK). A 40-mm (diameter) disk was compressed into the gel and redrawn. Velocity of the insertion assay was 1mm/s and withdrawn distance was 9mm. Adhesion was determined at 20°C and 37°C, and (iii) Microstructure of the hydrogel was assessed with Scanning Electron Microscopy (SEM).

#### Gelation assay

was carried out to determine the behavior of the hydrogel under UV light radiation at different temperatures. The following scenarios were tested: (i) 1mL of hydrogel and 15 μL of photo-initiator solution were mixed and placed on a petri dish. The hydrogel was heated at 37°C for 30 min and gelation was tested. Next, the hydrogel was cooled. (ii) 1 mL of hydrogel and 15 μL of photo-initiator solution were mixed and placed on a petri dish. The dish was placed under ultraviolet light 2000 mV/cm2 for 3 minutes (AnalytikJeta UVP Crosslinker) keeping the temperature below 20°C. The hydrogel was heated at 37°C for 30 min and gelation was tested. Next, the hydrogel was cooled. (iii) 1mL of hydrogel was placed on a petri dish. The hydrogel was heated at 37°C for 30 min and gelation was tested. Next, the hydrogel was cooled. (iv) 1 mL of hydrogel was placed on a petri dish. The dish was placed under ultraviolet light 2000 mV/cm2 for 3 minutes, keeping the temperature below 20°C. The hydrogel was heated at 37°C for 30 min and gelation was tested. Next, the hydrogel was cooled. For hydrogel observation after the irreversible episode, they were lyophilized at 50ºC for 96h and were examined by Field-Emission Scanning Electron Microscopy (FE-SEM) on a FEI Quanta 650 microscope using aluminum tape as support.

#### Degradation assay

was performed to understand the degradation rate of our hydrogel under different conditions that would try to mimic the different conditions found in the gastrointestinal tract, 1 mL of hydrogel was irreversibly gelified and then, submerge in 10mL of one of the following solutions: PBS pH=7, PBS pH=3, PBS pH=9, PBS pH=11 and rat feces bacteria culture on Luria-Bertani (LB), broth at a starting range of 105 CFU/mL. The piece of gelified hydrogel was weighed before the submersion (Wo) and then on set date points (Wt) until complete dissolution of the hydrogel or until one month under the conditions was reached. Each time point the medium was also changed when the weighting took place.

#### Drug absorbance and drug release kinetics

of the hydrogel was also evaluated *in vitro* with mathematical models to study the capacity of our hydrogel to both release and absorb molecules of different molecular weight, in order to further study the capability of this hydrogel to act as a drug eluting platform. For these experiments, we worked with two molecules: (i) Trypan Blue, which is a dye used in Biology as a vital stain to selectively color dead tissues or cells blue. Its chemical formula is C34H24N6Na4O14S4 and has a molar mass of approximately 960 Daltons (ii) Bovine serum albumin (BSA) which has a molecular weight of 66.5 kDa. For the Trypan blue experiments, first, a calibration curve was prepared by measuring the absorbance (λ=450 nm) of different concentrations of trypan blue diluted in water using a Uv-Vis spectrophotometer (Varioskan Flash, Thermo) SkanIt Software 2.4.1 was used to analyze results. Then, the curve equation was obtained: y=6234.9x+0.1008. R² = 0.99349. Data not shown. Absorbance and release of BSA was evaluated with the Bicinchoninic acid (BCA) Protein Assay Kit (PierceTM, Thermo Scientific).

The BCA Protein Assay Kit is a colorimetric detection and quantification method of total protein. It is based on the reduction of Cu+2 to Cu+1 by proteins in an alkaline medium (Biuret reaction) with the detection of cuprous cation (Cu+1) using BCA. The purple-colored reaction product of this assay is formed by the chelation of two molecules of BCA with one cuprous ion. Plates are then read at 562nm using a spectrophotometer. SkanIt Software 2.4.1 was used to analyze results.

To incorporate substances inside the hydrogel, 0.25 mL of gelated hydrogel were submerged in 5 mL of 0.0005 mg/mL Trypan Blue or 2 mg/mL BSA. Hydrogels incorporated the solution at room temperature for 48 hours. To calculate the amount of substance incorporated into the matrix of the hydrogel, the following equation was used: Incorporating efficiency (%): [(V_1_C_1_ - V_2_C_2_)/ V_1_C_1_] X100. Where: V1: original volume of substance solution (mL). V2: remaining volume of substance solution after 48h (mL). C1: initial concentration of substance (μg/mL) and C2: remaining concentration of substance (μg/mL).

Hydrogels were withdrawn from the solution and placed in release medium (PBS). At specific time points, amount of substance released into the medium was evaluated as mentioned before. The kinetics of drug release from the hydrogels was studied with three different models to asses which one fitted best with the release data from our experiments. The three different models were: Higuchi release model: M_t_ = K_h_t^1/2^, Zero - order release model: M_t_ = K_0_t, and First - order release model: M_t_ = 1 - e ^−K_1_t^. Where Mt is the fraction of drug released at each time point (t) and Kh, K0 and K1 are the Higuchi release kinetic constant, the zero-order release kinetic constant and the first-order release kinetic constant respectively.

Lastly, the drug release mechanism was analyzed using the Korsmeyer-Peppas model [F = (M_t_/M) = K_m_t^n^]. This is a semi-empirical model used when the exact mechanism is not known or in the case that more than one mechanism is involved in the drug release. Where F is the fraction of drug released at time (t), Mt is the amount of drug released at time (t), M is the total amount of drug in the hydrogel, Km is a kinetic constant and n is the release exponent, indicative of the drug release mechanism. n is estimated from linear regression of log (Mt/M) versus log t; when determining the n exponent, only portions of the release curve ≤ 60% should be used.

#### Biocompatibility assays (cytotoxicity, hemolysis and acute toxicity)

were first performed to check the possible cytotoxicity induced by CoverGel, carrying out a co-culture of a cell line with different conditions involving our hydrogel. The cell line chosen for these experiments was Caco-2 (ATCC® HTB-37TM) a human colorectal adenocarcinoma cell line. For cytotoxicity evaluation, 3×104 cells/cm^2^ cells were cultured in 24-well cell plates under the following conditions: Normal culture medium (DMEM medium supplemented with 10% Fetal Bovine serum, 1% L-Glutamine and 1% Penicillin/Streptomycin) as a positive control. Normal culture medium supplemented with 10% Dimethyl sulfoxide (DMSO) as a negative control. Normal culture medium supplemented with 1%-2%-5%-10%-15% of hydrogel v/v. Cells were cultured at 37ºC and 5% CO_2_ for 24, 48, 72 and 96 hours. At specific time points cell viability was evaluated using Resazurin Sodium Salt (Sigma-Aldrich). Resazurin sodium salt (7-Hydroxy-3H-phenoxazin-3-one-10-oxide sodium salt) is a blue dye weakly fluorescent until it is irreversibly reduced by living organism to Resofurin, a highly red fluorescent. The reduction of Resazurin Sodium Salt correlates with the number of living cells. Samples are exposed to Resazurin Sodium Salts for 4 hours and then absorbance at 570 nm is analyzed using a spectrophotometer (Varioskan Flash, Thermo).

Hemolysis test was done *in vitro*. Two mL of gelified hydrogel were placed in 20 mL of sterile saline and incubated for 72h at 37ºC. After, the solution was collected and filtered with a 0.22 µm membrane. Two mL blood samples were freshly collected from six male Sprague Dawley rats into an anticoagulant tube and gently mixed, blood was then diluted with 2.5 mL of saline. For the hemolysis test 10 mL of the hydrogel extraction, distilled water, and saline were poured in 50 mL test tubes and placed at 37ºC for 30 min. After that, 0.2 mL of diluted blood were added to each test tube with gently shaking and the solution was placed at 37ºC for 1 hour. The absorbance of the samples after 1 hour was evaluated with a spectrophotometer (Varioskan Flash, Thermo) at 545nm. SkanIt Software 2.4.1 was used to analyze results. Hemolysis ratio (%) was calculated with the following formula: [A hydrogel extraction-A saline solution / A distilled water - A saline solution] x 100.

Acute toxicity of CoverGel was evaluated in nine male Sprague Dawley rats, 7 months of age (400-550 g). All experiments were approved by the ethics committee of the animal facility. Animals were divided in four groups: three subjects and a control group. Rats from the subject groups were injected with 10 mL/kg of our hydrogel into the abdominal cavity and rats from the control group were injected with 10 mL/kg saline into the abdominal cavity. Animals were observed daily after administration, evaluation of general conditions (activity, hair, feces, behavior and other clinical signs), body weight and mortality were done. Animals were euthanized at 3, and 7 days with anesthetic overdose and major organs (heart, liver, spleen, lung and kidney) were extracted, evaluated, and fixed in 10% formaldehyde solution. Organs were then stained with hematoxylin-eosin for histopathological evaluation. Blood was also extracted, and hematological and biochemistry analysis were performed.

The effect of subcutaneous implantation of CoverGel was evaluated in six male Sprague Dawley rats, 7 months of age (420-500g). Rats maintained the subcutaneous implantation for 10 days. Under anesthesia, a 2cm incision was done on the dorsal back of the rat and 0.5g of gelified hydrogel were introduced subcutaneously. The incision was then sutured, and rats were allowed to recover with food and water ad libitum. After 10 days, rats were euthanized and the hydrogel was extracted, cut in two, one piece was placed in Carnoy solution (60% ethanol, 30% chloroform, 10% glacial acetic acid) for histological evaluation and the other piece was lyophilized (Christ Loc-1m, B.Braun) and evaluated using SEM.

### Endoscopy

All endoscopic evaluations were carried out using an Olympus Video bronchoscope EVIS EXERA II (BF-1T180) type endoscope with an outer diameter of 6.0 mm, working channel diameter of 3.0 mm. Room air was used for insufflation during the endoscopy. Endoscopic application of hydrogel was done by positioning the tip of the catheter over the mucosal lesions.

### Acute experimental colitis (EC) animal study

The EC animal model was induced by the rectal instillation of 0.6 mL of 3.5% TNBS solution in 50% ethanol (Day 0). The peak of colitis was established 3 days post administration (Day 3) and animals were examined to evaluate the affection and apply treatments. Forty male Sprague Dawley rats (Harlan Laboratories Models SL, Barcelona, Spain) weighting 250-300 g, which lost at least 8% of the original weight (Day 0) and showed a circumferentially affected area at day 3, were included in this study. Animals were randomized in 5 groups: (i) Sham group: healthy animals with no administration of TNBS. (ii) Control Group: animals with EC treated with 1mL of saline, through the endoscope. (iii) CoverGel group: animals with EC treated with CoverGel without active substance. (iv) CoverGel + Infliximab (IFX) group: animals with EC treated with CoverGel with IFX, 1mg/mL. (v) CoverGel + Vedolizumab (VDZ) group: animals with EC treated with CoverGel with VDZ, 1mg/mL.

Animals were weighted and underwent colonoscopy on days 0, 3 and 7. Blood was extracted by a small cut on the tail, and serum was kept frozen for future evaluations. Ponderal evolution was studied as variation (Δ) of original weight at day 0 for each rat. Four days after treatment (Day 7) animal were euthanatized by an anesthetic overdose and liver and colon were evaluated. Ulcer site samples were cut and placed in formaldehyde for H/E staining and blinded histopathological evaluation. Histological study evaluated the maintenance of the intestinal architecture and the inflammatory cell infiltrate in a 0-3 scale for both characteristics and a final 0-6 scale. The scoring system (Table 1) followed was extracted from Ulrike Erben et al^9^. The appearance of colony forming units in liver homogenate was evaluated to establish bacterial translocation (BT) to the liver as a marker of a weak intestinal barrier due to inflammation.

**Table 1.** Histological scoring system for the experimental colitis animal models.

| Inflammatory cell infiltrate: |  |  | Intestinal architecture: |  |  |
| --- | --- | --- | --- | --- | --- |
| Severity | Extent | Score 1 | Epithelial changes | Mucosal architecture | Score 2 |
| Mild | Mucosa | 1 | Focal erosions |  | 1 |
| Moderate | Mucosa and submucosa | 2 | Erosion | ± Focal ulcerations | 2 |
| Marked | Transmural | 3 |  | Extended ulcerations ± granulation tissue ± pseudopolyps | 3 |
| Sum of scores 1 and 2: |  |  |  |  | 0-6 |

### Chronic EC animal study

All animals included in this protocol underwent 4 rounds of TNBS colitis induction in the same fashion as the acute model, thus, induction was done on days 0, 7, 14 and 21, treatments were given on days 3, 10, 17 and 24 and euthanasia was performed on day 28. Twelve male Sprague Dawley rats were included. Animals were randomized in two groups: (i) IFX s.c. group: animals with EC treated with 5 mg/kg of IFX subcutaneously. (ii) CoverGel + IFX group: animals with EC treated with CoveGel+ IFX (1 mg/mL). All assessments were carried out the same way as in the acute EC study. Animals underwent endoscopic follow-up at days of induction, days of treatment and day of euthanasia. Furthermore, antibodies to IFX (ATIs) in serum were also evaluated. To separate ATIs from IFX as the might be found on blood, a protein G column and acidic buffer treatment were used. 600 µL of a 1:10 dilution of serum was applied onto the protein G column (Ab Spin trap; GE Healthcare) and washed. Trapped ATIs were eluted with 400 µL of elution buffer (0.1M Glycine-HCl pH 2.7) which releases proteins trapped in the protein G column and at the same time, dissociates ATI’s-IFX complexes. Samples were eluted into neutralizing buffer (1M Tris-HCl pH 9) containing an IgG concentration of 20 µg/mL to prevent the re-formation of immune complexes. Then, with the samples obtained from the protein G column, a 96-well plate was coated with 200 µL/well O/N at 4ºC or for 1 hour at 37 ºC. Plates were washed with washing buffer (PBS 0.1% Triton) and 200 µL/well Horseradish peroxidase conjugated-Infliximab (HRP conjugation Kit, abcam) (0.2 µg/mL IFX-HRP) were added for 30 minutes at room temperature, protecting the plate from direct light. The plate was further washed and finally, a substrate solution (3,3’,5,5’-Tetramethylbenzidine/H2O2 50% vol/vol) was added to the microplate. The enzyme reaction yields a blue product that turns yellow when stop solution (2 M sulfuric acid) is added. The intensity of the color measured is in proportion of the quantity of ATIs bound in the initial step; this intensity is measured by determining the optical density of each well using a microplate reader at 450 nm with a wavelength correction set to 540 nm using a spectrophotometer (Varioskan Flash, Thermo). SkanIt Software 2.4.1 was used to analyze results.

### Statistical study

All values are expressed in median ± range unless otherwise stated. Comparison of groups was done using 2 ways ANOVA and Tuckey’s multiple comparison tests as a posthoc. Release kinetics was studied using mean values of 6 different experiments and a nonlinear regression fit to each different model equation. Graphpad Prism 6.0 software was used to perform analysis.

## RESULTS

### Development of the drug-eluting platform

All hydrogels showed a similar adhesion force at 22ºC, but when temperature was raised, adhesion force of only CD2 (CoverGel) was highly increased, from −25 to −3993 mN/s (Figure 1A). Viscosity (η) of all samples was similar at 22°C, but when temperature was increased to 37°C, viscosity of CD2 was increased from <1 Pa⋅s at 22 ºC to 1500 Pa⋅s at 37 ºC (Figure 1B). In addition, the morphology of irreversible hydrogel was evaluated by SEM observing that it is formed by a structured network with well-defined porous distribution ranging from 50-300 μm (Figure 1C).

**Figure 1.**
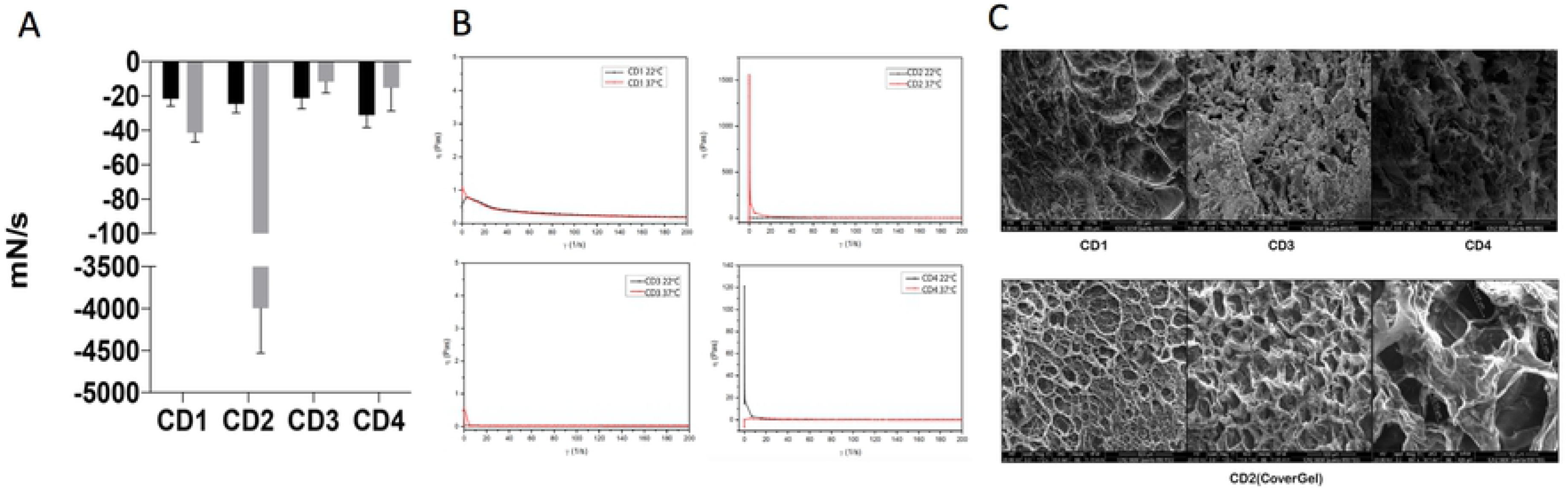
Rheological evaluations of the four original colon dressings (CDs). (A) Adhesion force evaluated at 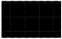 22°C and 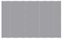37°C expressed in mN/s. (B) Viscosity evaluated at 22°C (black line) and 37°C (red line) expressed in pascals. (C) Scanning electron microscopy images of CD1, CD3 and CD4 (top three pictures) and CD2 (bottom pictures).

Reversible and irreversible gelation of the hydrogel was acquired. Hydrogel can be given smoothly through a catheter (diameter 2.0–2.2 mm) by applying a force of 370 mmHg (0.48 atmospheres). Hydrogel was also stable during 30 days at physiological-like conditions (Figure 2A). For Trypan Blue, the incorporating efficacy was 58.96 ± 2.8% and for BSA 24.09 ± 4.34% (mean ± SD). Drug release from hydrogel was related to the concentration of the drug in the medium, which followed a Fickian diffusion. The release kinetics model that best fits our results is the Higuchi release model, being the final equation obtained for our results: for Trypan Blue (Mt=0.05760t^1/2^; R^2^=0.8986) and for BSA (Mt=0.1369t^1/2^; R^2^=0.9167). The release exponents (n) were 0.4338 for Trypan Blue and 0.5116 for BSA. (Figure 2B). Table 2 shows correlation coefficient of the different kinetic models and the Korsmeyer-Peppas mechanism model.

**Table 2.** Coefficient of determination (R^2^) of our data to the three kinetic release models (zero-order release model, first-order release model and Higuchi release model) and for the drug release mechanism (Korsmeyer-Peppas model).

| Sample | 0-order $R^2$ | 1<sup>st</sup>-order $R^2$ | Higuchi $R^2$ | Korsmeyer-Peppas |
| --- | --- | --- | --- | --- |
| Tryp. B | 0.2022 | 0.3361 | 0.8986 | 0.9169 |
| BSA | 0.5393 | 0.8324 | 0.9167 | 0.9171 |

**Figure 2.**
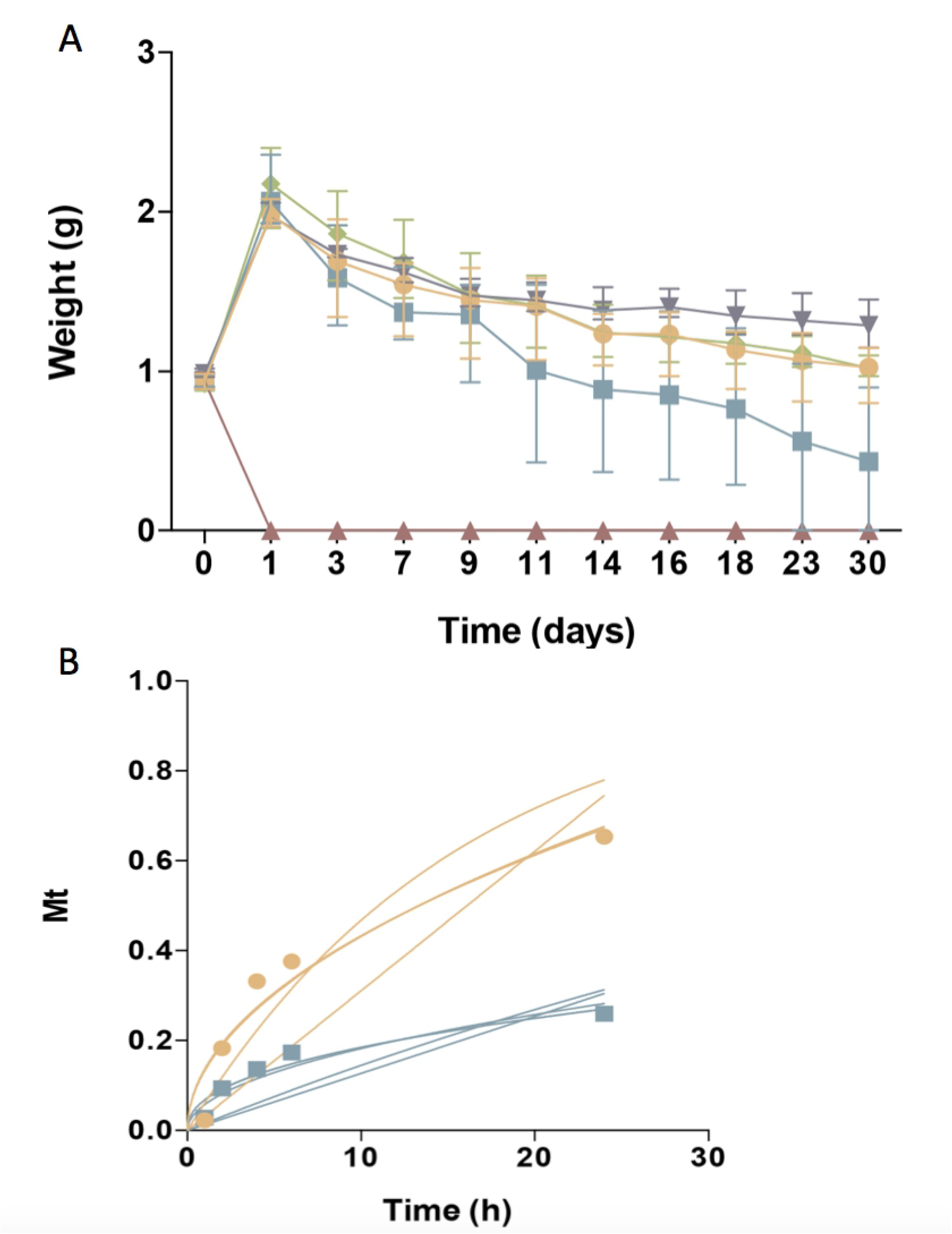
Characterization of CoverGel. (A) Degradation of the hydrogel when placed for 30 days under different conditions (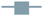 PBS pH3, 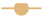 PBS pH7, 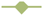 PBS pH9, 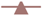 PBS pH 11 and 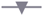 bacterial culture). Graph represents weight of hydrogel over days. (B) Drug-release kinetics for 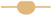 Bovine serum albumin and 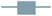 Trypan blue. Graph represents concentration of substance on release medium (Mt) over time. Lines resepren the different mathematical models tested.

CoverGel showed good biocompatibility, both *in vitro* and *in vivo* studies. Caco-2 cells cultured with different concentrations of our hydrogel showed an increased proliferation, reaching a top at 10% when compared to the positive control of the experiment (normal culture medium) (Figure 3). Hemolysis rate caused by our hydrogel on blood from three different rats was 2.829 ± 1.135%.

**Figure 3.**
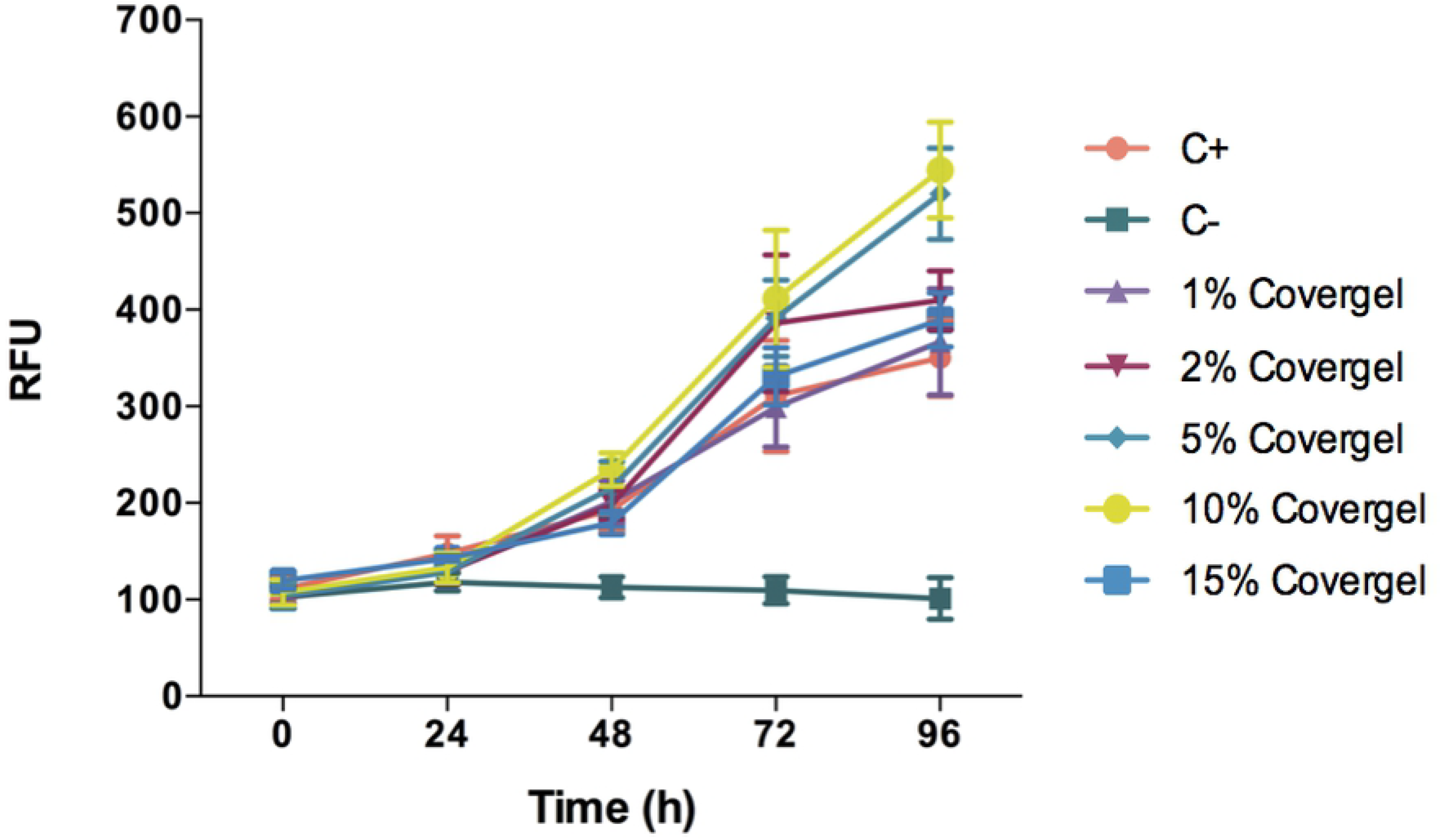
Cell viability of Caco-2 cells in conditioned medium with 1%, 2%, 5%, 10% and 15% of CoverGel. Cells were cultured at 37ºC and 5% CO_2_ for 24, 48, 72 and 96 hours. Values are expressed in Relative Fluorescence Units (RFU).

There was no sign of inflammation, toxicity or tissue malformation on major organs induced by the hydrogel. Macroscopic evaluation of major organs didn’t show any signs of abnormalities, and organs weight with respect of total body mass didn’t present any significant differences (Figure 4A). Heart tissue didn’t show signs of toxicity, cardiac myocytes maintain a good arrangement, and no hemorrhage, inflammation nor necrosis is observed. In the liver, no hepatocellular degeneration occurs, and there isn’t an abnormal neutrophil, lymphocyte or macrophage infiltrate in the tissue. Tissue structure from spleen, kidneys and lungs is maintained as well (Figure 4B). Subcutaneous placement of 0.5g of the hydrogel showed a synovial metaplasia, a capsular surface covered by fibrohistiocytic cells that are grouped forming pseudovilli. No toxicity or inflammation was seen in the surrounding tissue, meaning that the body responded by encapsulating the hydrogel as it would do with a foreign body, but no rejection response was triggered (Figure 4C). At the time of placement, hydrogels weighted 0.5 g; when extracted, the weight was 0.95 ± 0.06 g. The platform was able to maintain its porosity and inner structure. No cell infiltration was seen 10 days after subcutaneous placement. (Figure 4D).

**Figure 4.**
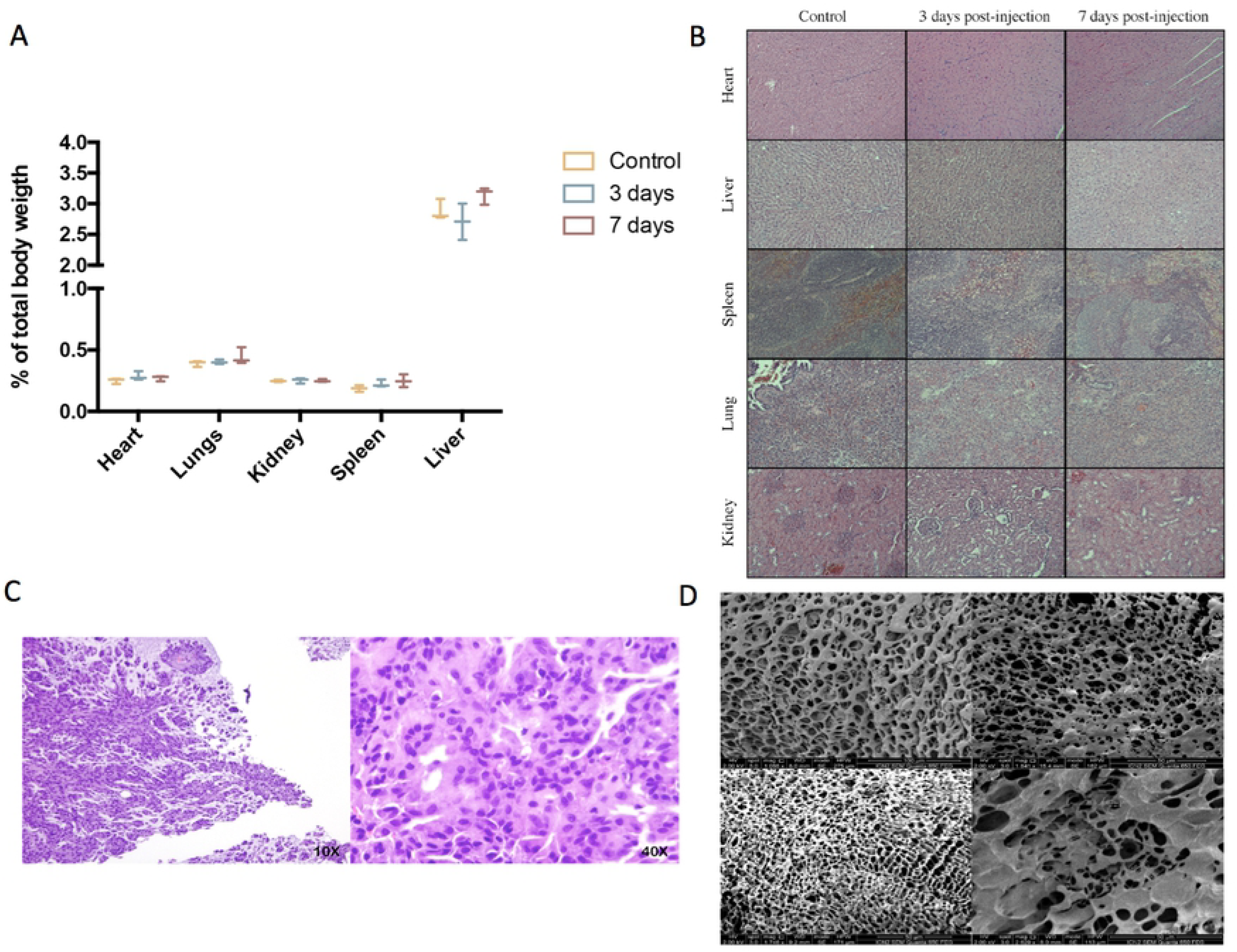
*in vivo* biocompatibility study. (A) % of organ weight relative to total body weight after intraperitoneal injection of 10ml/kg of CoverGel on rats. 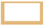 represents control rats 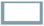 rats euthanized three days post injection and 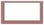 rats euthanized 7 days post injection. (B) representative images of major organs H/E stainings from each group at a 40X zoom. (C) H/E staining images of tissue surrounding subcutaneouslly placed CoverGel after 10 days post placement (10X left image, 40X right image). (D) scanning electron microscopy images of hydrogel after 10 days post subcutaneous placement.

### Efficacy of the drug-eluting platform in acute EC animal study

At day 3 all animals presented an entire circumferential affectation of the wall. Macroscopic and endoscopic evaluations at day 7, showed a complete mucosal restoration (vascular pattern, ulceration and friability) in CoverGel+IFX and CoverGel+VDZ groups, whereas in CoverGel group, this mucosal restoration was not complete. However, all treated groups clearly showed less mucosal damage than Control group (Figure 5). All animals with acute EC had lost more than 8% of original weight at day 3. At day 7 control animals showed a significantly weight loss (−55%), which confirm severe disease, as compared with the other groups. Drug-eluting platform with IFX or VDZ showed a significantly weight recovery in comparison with CoverGel without bioactive drug (−17% and −20% vs. −36%; p<0.044) (Figure 6A).

**Figure 5.**
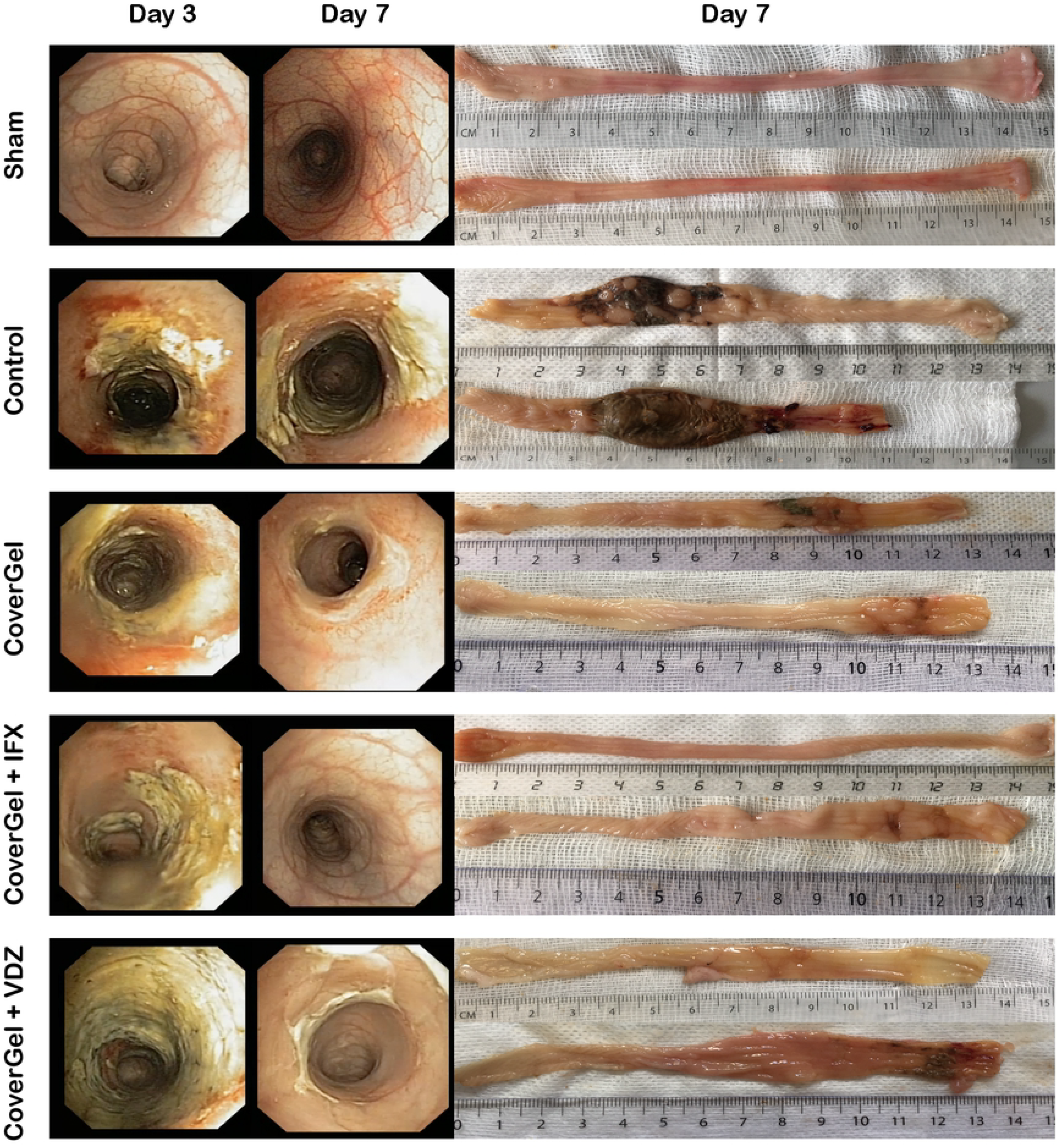
Endoscopic and macroscopic representative images of the acute EC animal model. Endoscopic images from day 3 (day of treatment) and day 7 (day of euthanasia) and macroscopic image of day 7 (day of euthanasia) from two representative animals of each group.

**Figure 6.**
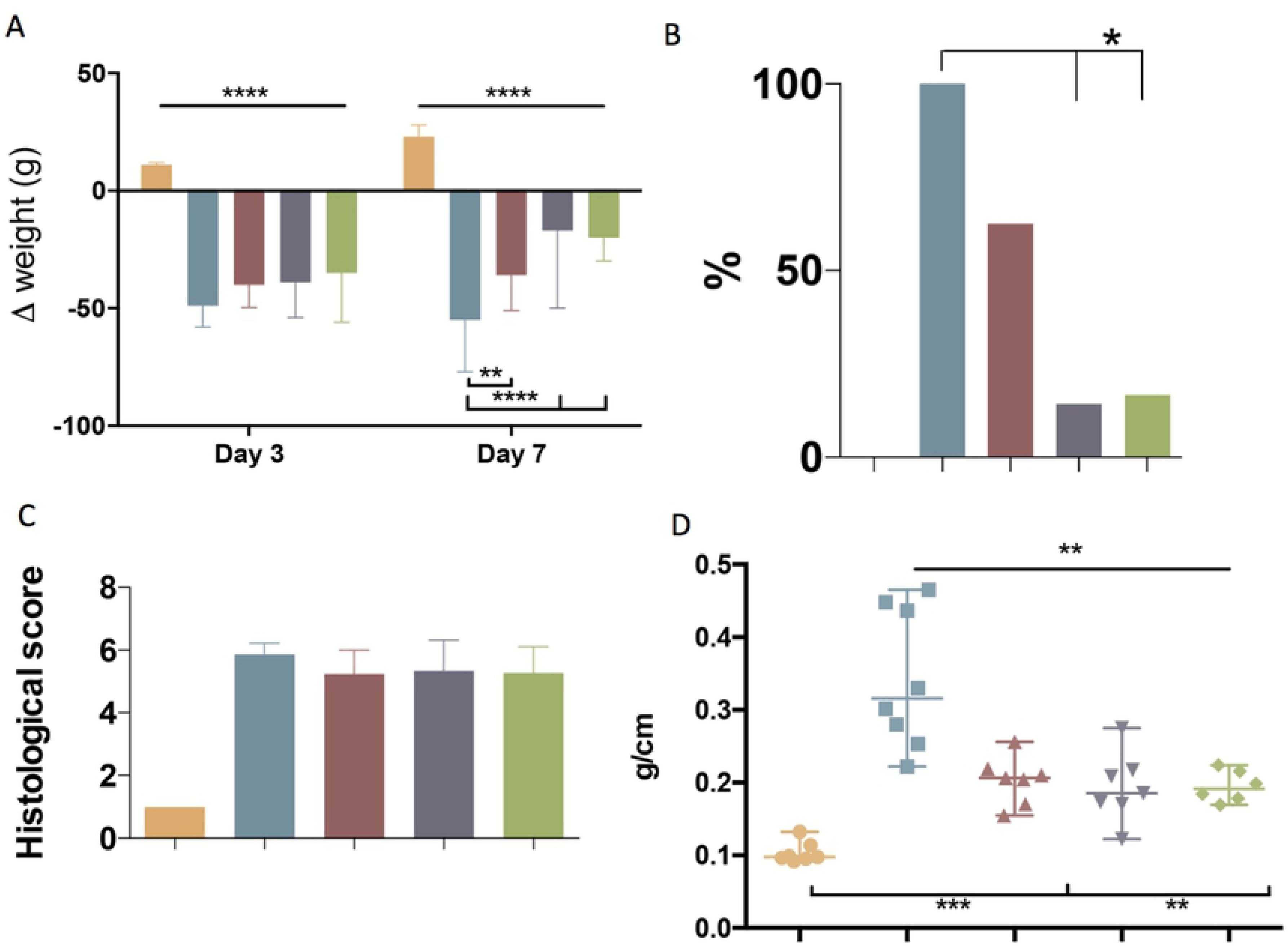
Clinical evaluations on rats from the acute EC animal model. Color representations are as follow: 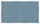 Control group 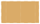Sham group 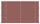CoverGel group 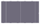 CoverGel + IFX group and 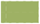CoverGel + VDZ group. (A) variation (Δ) of weight relative to day 0 (day of induction. (B) bacterial translocation to the liver. (C) histological score. (D) Colon weight/length ratio.

BT to liver at day 7 was absent in sham animals. Control group presented a significantly higher BT as compared with CoverGel+IFX and CoverGel+VDZ (100% vs. 14.3% and 16.6%; p<0.05). Animals treated with CoverGel showed a decreased BT (62.5%) without statistical differences (Figure 6B). Histological score for groups with EC was comparable: Control 5.87 ± 0.35, CoverGel 5.24 ± 0.76, CoverGel+IFX 5.34 ± 0.98 and CoverGel+VDZ 5.28 ± 0.82. (Figure 6C). The ratio between colon’s length and weight (Median (Range)), a simple indicator of edema and inflammation in the tissue, was significantly higher in control animals as compared with the other treatment groups (Control vs. CoverGel, CoverGel+IFX and CoverGel+VDZ) (0.3157 (0.244) g/cm vs. 0.2063 (0.1012) g/cm p=0.0012, 0.1854 (0.1527) g/cm p=0.0012 and 0.1914 (0.0545) g/cm p=0.0013), respectively (Figure 6D).

### Chronic EC animal study

There was variation in both groups in the degree of colitis induction throughout the 4 TNBS applications. Figure 7 shows representative endoscopic and macroscopic images of both groups, showing ulcerated mucosa covered by feces despite lavages. No statistically significant differences were observed in ponderal evolution at any day of the study (Figure 8A), in BT to liver (two animals out of six in each group, 33.3%) (Figure 8B) and in histological score for CoverGel+IFX (5.48 ± 0.55) and IFX s.c. (4.96 ± 0.71) (Figure 8C). Colon weight/length ratio at day 28 was significantly lower in the IFX subcutaneous treated group compared to the group treated with CoverGel+IFX (0.213 (0.124) vs. 0.3497 (0.129), p=0.0087) (Figure 8D).

**Figure 7.**
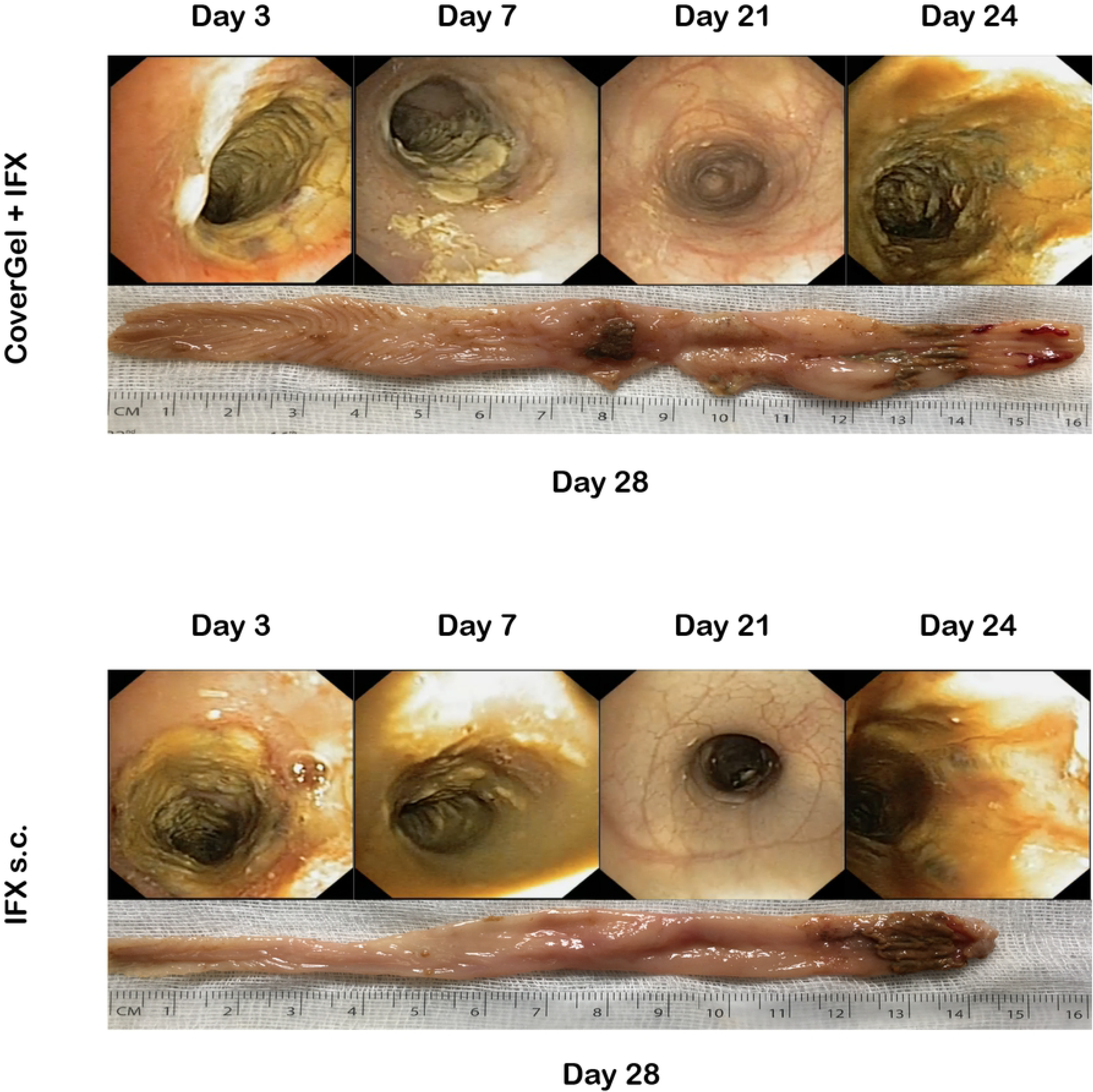
Endoscopic and macroscopic representative images of the chronic EC animal model. Endoscopic images from day 3 (first treatment application), day 7 (second TNBS induction), day 21 (fourth TNBS induction) day 24 (fourth treatment application) and macroscopic image of day 28 (day of euthanasia).

**Figure 8.**
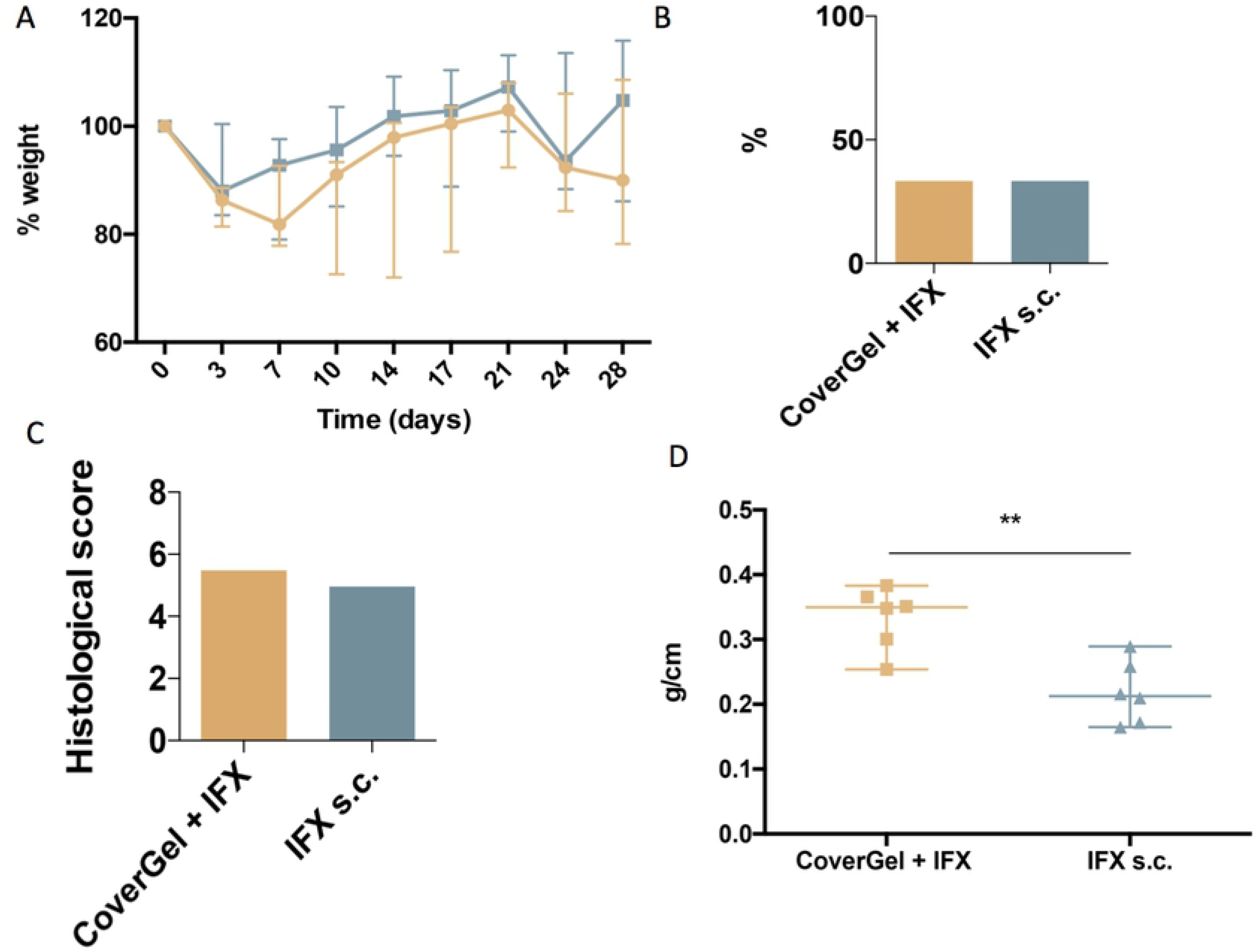
Clinical evaluations on rats from the chronic EC animal model. Color representations are as follow: 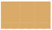CoverGel + IFX group and 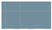IFX s.c. group. (A) variation (Δ) of weight relative to day 0 (day of induction). (B) bacterial translocation to the liver. (C) histological score. (D) Colon weight/length ratio.

The evolution of levels of ATIs for each group is shown in Figure 9. At day of euthanasia, levels of ATIs were significantly lower in rats treated with CoverGel+IFX compared to rats treated with IFX subcutaneously (0.90 ± 0.06 μg/mL-c vs. 1.97 ± 0.66 μg/mL-c, p=0.0025).

**Figure 9.**
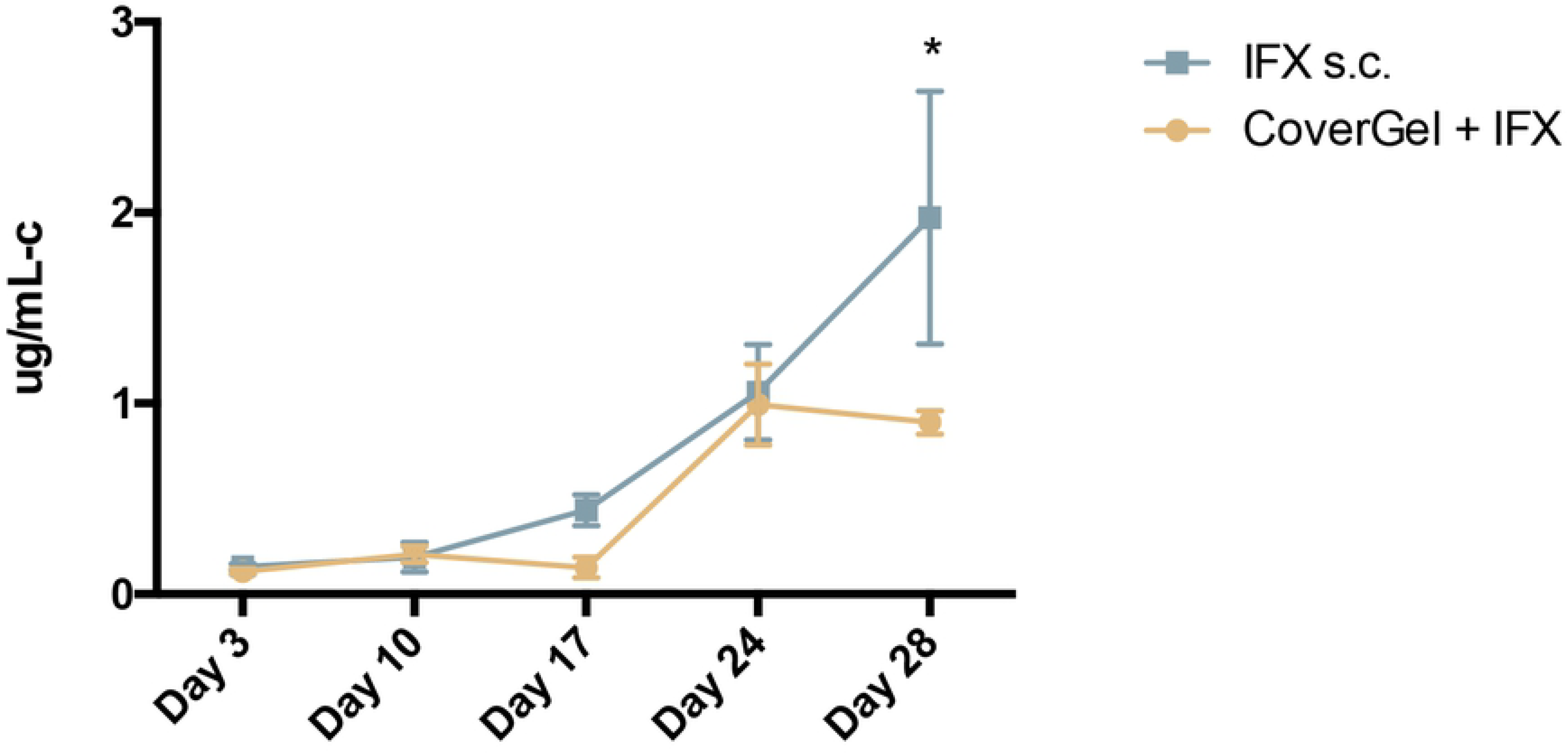
Evolution of antibodies to Infliximab (ATIs) in serum for: 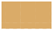 CoverGel + IFX group and 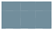 IFX s.c. group. Graph represents the concentration of ATIs (μg/ml-c calibrated in comparison to a polyclonal rabbit α-IgG standard).

## DISCUSSION

Mucosal lesions refractory to biological treatments in IBD patients is a clinical challenge and unmet need that requires new treatment modalities. Drug delivery directly to the colon is an option thanks to oral dosage formulations that are designed to achieve a controlled drug release. However, this controlled release has mainly been achieved with small molecules but not for macromolecules, such as biologics^10^. Direct release in the colon is beneficial to prevent early absorption of the drug, thus increasing the quantity of drug reaching the target tissue and at the same time decreasing systemic exposure and associated side effect. A specific form of control drug release is polymer-based drug delivery. Polymers offer a wide range of possibilities useful for colonic delivery. These polymers can be triggered to start releasing drug by physiological characteristics of the healthy colon, as well as in different disease situations^10^. Hydrogels for drug delivery have also been studied for their capacity of mucoadhesion and controlled delivery^11^.

Local injection of infliximab can be a novel scenario for endoscopic therapy in IBD patients with symptomatic isolated mucosal lesions, as has been proposed by our group^12^. The real efficacy of locally injected IFX is not discernible from current literature, but available data seem encouraging at dose of 100 mg per session, assuming that higher anti-TNF concentrations given through local injection can favor healing and prevent adverse events, because a trend for a dose–response curve was observed. Topical injection of anti-TNF agents has been described in the treatment of long-standing perianal fistulizing CD in the absence of abscess^2^. There are technical variations in topical injection of the anti-TNF agents. The agents were injected in a circumferential or intrafistula fashion via the external or internal.

In this work we have provided new therapeutic tool for endoscopy by developing a new hydrogel (CoverGel) based on HA, MC and F127 which presents biocompatible, bioadhesive and bioactive properties^13–17^. CoverGel is an endoscopic vehicle to locally deliver biological drugs (IFX and VDZ) with proven efficacy in acute and chronic EC in rats. Moreover, endoscopic administration of IFX induces significantly lower levels of ATI as compared with subcutaneous administration. The rationale for the drugs dosages used (1 mg/mL) with our drug eluting gel are based on previous clinical experience with local injection of IFX^12^.

Our developed hydrogel showed a reverse thermo sensitive capacity of gelation accompanied by a greater adhesion capacity. When the hydrogel contacts the mucosal layer at a physiological temperature, it gellifies permitting the adhesion to the mucosa^18^. Rheological studies in our hydrogel confirmed that it is a non-Newtonian fluid, meaning that its viscosity varies with temperature. Morphologically, our hydrogel presents a macroporous (pore diameters of greater than 50 nm) interconnected network that can be the matrix for cell ingrowth as well as nutrients and waste flow, essential characteristics for cell survival within the hydrogel. This platform is less vulnerable to matrix transformations due to osmotic pressure or tissue extrusion, which is an advantage for a controlled drug release of the hydrogel^19^. Different kinetic models were studied with the results obtained from the drug release experiments. The kinetic model that best fitted our results was the Higuchi release model. This model is explained by Fick’s law, in which diffusion of a molecule is related to its concentration. The diffusion flux goes from a higher concentrated area to a lesser-concentrated one. The magnitude of the diffusion flux will be proportional to the concentration gradient. These results and kinetic model were confirmed when mechanistic of drug release from Covergel were studied by the Korsmeyer-Peppas model, confirming that the diffusion coefficient follows a Fickian diffusion. Our hydrogel showed the capacity to improve cellular viability in a dose-dependent manner at least until a 10%, demonstrating a good potential to become a cell carrier. Intraperitoneal injection stated that CoverGel did not cause an acute toxicity reaction on animals three or seven days post intraperitoneal injection. Subcutaneous placement of our hydrogel in rats was done to evaluate *in vivo* behavior and biodegradability of our hydrogel, which was evaluated by SEM and we confirmed that the inner structure was maintained 10 days after s.c. placement and no cell infiltration were seen.

All of these results help us conclude that this hydrogel is a good candidate to be used in the digestive system and implement it in endoscopic techniques in order to cover some unmet needs of the field. Although in vitro and in vivo results are really promising, further experiments on in vivo drug release kinetics and degradation need to be performed, as these characteristics have previously been reported to vary from in vitro analysis to in vivo, due to inflammatory cell response^20–22^.

TNBS induces colitis has been proven as a valid method to elucidate different mechanisms underlying the mechanism of the disease^23^. Furthermore, and besides its limitations mimicking population refractory to biological treatments, this model is a great tool to study the immunopathogenesis of the disease and the potential benefit of different therapies^23^.

Our results show that the simple application of our hydrogel as a dressing on the inflamed tissue by means of a shield to improves the overall outcome and restoration of the mucosal layer. This beneficial effect of CoverGel is based on its good biocompatibility properties, able to improve mucosal healing. However, when CoverGel is loaded with biological drugs with proven efficacy treating this disease, the effect is even more pronounced. In this sense, endoscopic response was obtained after 4 days of treatment in all animals treated with CoverGel, but unfortunately histological remission was not achieved at this time. Despite we do not have measurements of local and systemic levels of the drugs applied by the hydrogel, clinical response was accompanied by different clinical aspects such as body weight loss, colon weight/length ratio or bacterial translocation to the liver, all features improved in the groups treated with CoverGel alone or in combination with drugs.

The efficacy of an enema based on ascorbyl palmitate hydrogel loaded with dexamethasone has been demonstrated in mice with colitis as well as in *ex vivo* analysis of colon samples from patients with IBD^24^. By contrast, our hydrogel, which can be also administered as enema, has been designed to be administered under endoscopic view to perform a more localized treatment. Another goal of our project was to improve the patient developing resistance to anti-TNF agents, which is one of the main problems associated with biologics. This resistance is caused by the formation of antidrug antibodies (ADAs), as a response of the body to these exogenous complex proteins. The kind of mAb’s most used in IBD, no matter their nature, cause to a greater or lesser degree ADAs formation^25^. Since new drug development is a complex and long-lasting effort, new therapeutic approaches are needed to reduce this risk, and allow patients to continue on a successful therapy longer. The direct drug administration with CoverGel was able to significantly improve the disease outcome in an animal model using a much lower dose than the standard dose applied. Moreover, when we compared the formation of ATI’s from subcutaneous injection vs. CoverGel application in an experimental model of chronic EC (with a longer period of time of the disease state) we observed that whereas there were no major significant differences on the outcome of the disease between both ways of treatment application, the levels of ATI’s were significantly lower on the group treated with CoverGel. These results confirmed our theory that a locally applied therapy at a lower dose is able to obtain a good clinical result and at the same time reduce adverse events associated with the systemic application of a drug.

Taken together, results from our study, although preliminary and preclinical, demonstrate that endoscopic administration of a based-on hydrogel drug delivery platform represents a safe and promising approach for the management of active lesions with a lower dose as compared with systemic administration. Present findings also suggest than levels of ADAs are lower with this approach, opening the opportunity to be used in patients with past history of acute infusion reactions to the drug. Further research, preclinical and clinical, needs to be pursued to safely evaluate and introduce this technique in standard practice to increase the pool of therapeutics interventions to treat IBD patients. Direct drug administration through endoscopy opens new possibilities for therapeutic endoscopy, allowing for the application of bioactive treatments in different pathologies already managed with endoscopy such as colorectal cancer and local inflammatory lesions.

## Brief summary

Direct drug administration through endoscopy opens new possibilities for therapeutic endoscopy in IBD. CoverGel is an endoscopic vehicle to locally deliver biological drugs with proven efficacy in acute and chronic experimental colitis in rats and inducing less immunogenicity reaction.

## Author Contributions

Conceived and designed the experiments: IB, RB, VL-Z. Performed the experiments: IB, MC-S, RB, VL-Z. Analyzed the data: IB, NO. Contributed reagents/materials/analysis tools: RB, VL-Z, MC-S. Wrote the paper: IB, RB, VL-Z.

